# *Mycobacterium tuberculosis* sulfolipid-1 (Sl-1) increases the excitability of mouse and human TRPV1-positive sensory neurons in a YM254890-reversible fashion

**DOI:** 10.1101/2025.06.28.662105

**Authors:** Dhananjay K. Naik, Felipe Espinosa, Ifunanya M. Okolie, Kubra F. Naqvi, Giaochau Nguyen, Cody R. Ruhl, Sven Kroener, Gregory Dussor, Michael U. Shiloh, Theodore J. Price

## Abstract

Cough is a hallmark sign of tuberculosis and key driver of transmission. While traditionally attributed to host-driven inflammation, we previously demonstrated that *Mycobacterium tuberculosis* lipid extract (Mtb extract) and its component sulfolipid-1 (SL-1) directly activate nociceptive neurons to induce cough in guinea pigs. However, the cellular mechanisms by which Mtb extract and SL-1 modulate nociceptive sensory neurons remain incompletely understood. Here, we show that Mtb extract enhances action potential (AP) generation in mouse nodose nociceptors via an SL-1–dependent mechanism. Using calcium imaging, we found that Mtb extract and SL-1 increased intracellular Ca²⁺ signals in TRPV1⁺ neurons from both mouse nodose and human dorsal root ganglia (hDRG). These Ca²⁺ signals were attenuated by the Gαq/11 pathway inhibitor YM254890, even in the absence of extracellular Ca²⁺, suggesting involvement of intracellular Ca²⁺ stores. Together, these findings indicate that SL-1 engages Gαq/11-coupled pathways to sensitize nociceptors via intracellular Ca^2+^ release, providing mechanistic insight into tuberculosis-associated cough and potential targets for therapeutic intervention.

## INTRODUCTION

Cough is a basic physiological process that results from irritation of the respiratory tract that serves to protect and help clear the airway (Bonvini and Belvisi, 2017; Brooks, 2011; Burnstock, 2006; Naqvi et al., 2023). Cough is triggered by activation of nociceptive sensory neurons that innervate the airway. The cell bodies for airway innervating nociceptors are located in both the nodose ganglion and thoracic dorsal root ganglion (DRG), and such nociceptors express canonical ion channels like transient receptor potential vanilloid 1 receptor (TRPV1) and Nav1.8 that are critical for the detection of and transduction of pain signals (Canning, 2011; Canning and Spina, 2009; Kim et al., 2022; Narula et al., 2014; Talbot et al., 2015). Cough is ultimately triggered by a central brainstem reflex activated by peripheral nociceptors (Canning, 2002; Canning, 2007; Canning et al., 2014; Footitt and Johnston, 2009; Ochoa-Cortes et al., 2010) (Canning, 2002, 2007; Canning et al., 2014; Footitt and Johnston, 2009; Ochoa-Cortes et al., 2010). Cough can be evoked in healthy individuals but also is a hallmark sign of respiratory infections (Burnstock et al., 2012; Canning, 2002; Canning, 2007a, 2008; Canning, 2011; Canning et al., 2014; Canning and Spina, 2009; Naqvi et al., 2023). Pathogens like Mycobacterium *tuberculosis* (Mtb) that are transmitted via the airborne route have evolved mechanisms that can engage pulmonary neuronal networks, triggering a cough (Garcia-Vilanova et al., 2019; Turner and Bothamley, 2015). Cough is one of the most prominent signs observed in individuals suffering from pulmonary tuberculosis (TB) (Bryant et al., 2016). According to the American College of Chest Physicians’ guidelines, a prolonged cough should trigger TB screening in individuals with epidemiologic risk factors (Field et al., 2018; Patterson and Wood, 2019). Cough, through the production of aerosols, has been implicated in the infectious nature of TB (Converse et al., 2003; Huddart et al., 2023; Jones-López et al., 2014; Loudon and Brown, 1967; Loudon and Spohn, 1969; Patterson and Wood, 2019; Perla, 1927; Seabra and Duarte, 2021; Singh et al., 2020).

The prevailing paradigm had been that Mtb triggers cough through its interaction with the host immune system, leading to irritation of the lung epithelium and the subsequent release of inflammatory mediators. However, we recently demonstrated that a total lipid extract from *M. tuberculosis* (Mtb extract) was sufficient to elicit coughing in guinea pigs (Ruhl et al., 2020). Moreover, Mtb extract was shown to directly activate mouse nodose and mouse and human DRG neurons by inducing Ca^2+^ influx, with this effect primarily attributed to the presence of sulfolipid-1 (SL-1), a major component of Mtb extract. Given the substantial public health burden imposed by Mtb infection, uncovering the molecular mechanisms by which SL-1 triggers the cough reflex would aid the development of targeted antitussive therapies. An unanswered question is how SL-1 influences the generation of action potentials in nodose neurons that innervate the upper airway. One of the major goals of this study was to understand if SL-1 produces changes in excitability, as measured by action potential generation in mouse nodose neurons using whole-cell patch clamp electrophysiology.

Metabotropic G protein-coupled receptors (GPCR) have been implicated in cough reflex initiation through their ability to modulate the activity of ion channels expressed by sensory neurons (Maher, 2009; Migliori et al., 2021; Naqvi et al., 2023; Shajahan et al., 2016). It is well established that certain classes of GPCRs can trigger the release of intracellular Ca^2+^ through the Gαq/11 pathway (Predescu et al., 2019; Trent et al., 2024). Given our previous findings demonstrating that SL-1 induces Ca^2+^ signaling in nodose and DRG neurons (Ruhl et al., 2020), we hypothesized that SL-1 triggers the release of intracellular Ca^2+^ via the Gαq/11 pathway and that this release can be inhibited by the Gαq/11 inhibitor YM254890 (YM). Our findings demonstrate that Mtb extract and SL-1 induce Ca^2+^ signaling in mouse nodose and in human DRG neurons in a manner that is dependent on intracellular Ca^2+^ stores and attenuated by YM. Our results give new insight into how Mtb acts directly on mouse and human sensory neurons, mechanisms that are likely important for how Mtb produces cough.

## METHODS

### Animal Studies

Experiments were done using Institute of Cancer Research (ICR, sometimes also referred to as CD-1) mice, aged 3-4 weeks, purchased from Envigo. Mice were maintained at the University of Texas at Dallas in a climate-controlled room on a 12:12 h light and dark cycle with food and water ad libitum. Animal procedures were approved by the Institutional Animal Care and Use Committees (IACUC) at the University of Texas at Dallas under protocol number 14-04 and were performed in accordance with the guidelines of the NIH Guide for the Care and Use of Laboratory Animals.

### Pharmacological tools

Three Mtb extracts were tested in this study: Wild type Mtb Erdman extract (Ruhl et al., 2020), Mtb Erdman Δ*stf0* extract lacking SL-1 due to genetic deletion of sulfotransferase 0 (*stf0*) (Converse et al., 2003; Mougous et al., 2004; Ruhl et al., 2020b) and Mtb Erdman Δ*stf0*::*stf0* extract, whereby the *stf0* gene was reintroduced at the attB site in Mtb Erdman Δ*stf0* (Mougous et al., 2004; Ruhl et al., 2020). Accumulation or absence of SL-1 was confirmed by mass spectrometry (Ruhl et al., 2020). Extracts were resuspended in dimethyl sulfoxide (DMSO) and diluted to a testing concentration of 0.4mg/ml based on prior studies (Ruhl et al., 2020). Isolated SL-1 was obtained from BEI resources as a purified molecule from Mtb (Manassas, Virginia, NR-14845). SL-1 was resuspended in DMSO and used at a concentration of 1 µM based on its EC_50_ of 33 nM (Ruhl et al., 2020b). YM, a known Gαq/11 GPCR inhibitor used in this study was purchased from Tocris biosciences (Bristol, UK,7352). YM was resuspended in DMSO and used at final concentration of 100 µM as used in previous studies (Dwyer et al., 2025; Peng and Shen, 2019; Trent et al., 2024). Capsaicin (Cap) was purchased from Millipore Sigma (St. Louis, MO,404-86-4) and dissolved in ethanol to create a stock of 1M and diluted to a concentration of 200 nM to test on mouse nodose and hDRG neurons (Tavares-Ferreira et al., 2022).

### Nodose ganglion cultures

Nodose ganglia were dissected bilaterally from male and female (12 males, 6 females) ICR mice and kept in a petri dish containing cold hanks’ balanced salt solution (HBSS, Gibco, Grand Island, NY, H6648) to remove the blood. Unwanted connective tissue was removed from the nodose tissue to clean it further. The tissue was transferred to a tube containing 5 mL of cold HBSS without Ca^2+^ and magnesium (Mg^2+^, Sigma, St. Louis, MO, H6648). The sample was centrifuged at 400 rotations per minute (rpm) for 2 minutes to facilitate the sedimentation of tissue fragments at the bottom of the tube. The tissue was then subjected to enzymatic digestion using collagenase A and collagenase D (each 1 mg/ml, Roche, Indianapolis, IN, 45-11088858001) with papain at 30 U/ml (Roche, 45-10108014001) for 20 minutes at 37 °C. After the digestion, the tissue was triturated in 1 ml of HBSS. The homogenized solution was then passed through a 70 µm cell strainer (Corning, NY, 352350) to remove debris. The filtrate was then gently added to a density gradient made from a 10% bovine serum albumin (BSA) (Biopharm, Bluffdale, UT, 71-015-025). The gradient was centrifuged at 900 x g (9 acceleration, 7 deceleration) for 5 minutes. The supernatant was then carefully removed, and the pellet was resuspended inDulbecco’s modified eagle’s medium (DMEM)/F12/GlutaMAX (Gibco, Grand Island, NY,10565018) culture media supplemented with 10% fetal bovine serum (FBS, Hyclone, Logan, UT, H30088.03) and 1% penicillin/streptomycin (Pen/Strep, Gibco, 15070-063). The cells were then plated onto dishes precoated with poly-d-lysine (PDL) (MatTek, Ashland, MA, P35GC-1.5-10-C) with laminin (Sigma-Aldrich, L2020). Cells were allowed to settle for 2 hours to adhere. After 2 hours, the cells were then supplemented with the same culture media with nerve growth factor (NGF, 10 ng/ml; Sigma, 01-125) and 5-fluoro-2’-deoxyuridine (FRD, 3 μg/ml, Millipore Sigma, F0503) + uridine (U, 7 μg/ml, Sigma-Aldrich, 856657) (FRD+U)) and incubated at 37 °C with 5% CO_2_ for 24 hours before use.

### Ca^2+^ Imaging with Mouse Nodose Cultures

After approximately 24 hours in culture, cells were loaded with Fura-2 AM (Ca^2+^ indicator dye, 1 ug/ml, Thermo-Fischer, Waltham, MA, F1221) for 1 hour, then transferred to a normal bath solution (135 mM NaCl, 5 mM HEPES, 1 mM CaCl_2_, 1 mM MgCl_2_, and 20 mM glucose). The pH and osmolality were adjusted to pH 7.4 and 300 mOsm with N-methyl-D-glucamine (NMDG, Millipore Sigma, Burlington, MA, M2004). Live cell Ca^2+^ imaging was performed using an Olympus IX73 microscope with MetaFluor Fluorescence Ratio imaging software. Cells were then imaged every second at 340/380 excitation wavelengths and 510 emission wavelength, and 340/380 ratiometric data were obtained for further analysis.

The cells on each dish were subjected to a 50-second baseline recording, followed by the addition of vehicle control (0.8% DMSO, Fischer Scientific, BP231-1) for 50 seconds. We then added the test compounds (Mtb extract, Mtb Δ*stf0* extract, SL-1 and/or vehicle for 150 seconds). For experiments involving YM (Tocris Biosciences, Bristol, UK, 7352) (with or without Ca^2+^ in the external bath), coverslips were pre-incubated in a bath solution containing 100 µM YM for 15 minutes before imaging. Subsequent imaging procedures were performed as previously described, with the continued presence of YM in the bath solution. Finally, Cap (Sigma, 404-86-4) at a concentration of 200 nM was used to determine if the cells express the TRPV1 (Caterina et al., 1997; Tominaga et al., 1998) which was applied at the end of the imaging session for 30 seconds. Positive responses were defined as a 20% change over the baseline.

For experiments with no external Ca^2+^, the external bath consisted of 135 mM NaCl, 5 mM KCl, 10 mM HEPES, 2 mM MgCl_2_, and 20 mM glucose. Cells were imaged using this external buffer as described above. At the end of the imaging session, Cap responses were elicited by replacing the external buffer with a Ca^2+^-containing solution and applying Cap for 30 seconds.

### Mouse Nodose Cultures for Electrophysiology

Nodose tissues were dissected bilaterally from ICR mice and transferred to a 5 mL tube containing cold HBSS. Cell cultures for patch clamp electrophysiology were prepared as previously described (Jeevakumar et al., 2020; Moy et al., 2017). The tissue was centrifuged at 400 rpm for 30 seconds to settle the dissected tissue. The cells were then incubated in 1 mL of papain 20 U/mL (Worthington, Columbus, OH 9001-73-4) for 15 minutes. Following this, the cells were treated in 1 mL of 3 U/mL of collagenase type II (Worthington, 9001-12-1) for 15 minutes. The digested tissue was then subjected to trituration through fire-polished glass pipettes with progressively smaller opening sizes. The triturated tissues were then centrifuged at 400 rpm for 5 minutes and plated on dishes coated with laminin and PDL. The plated cells were allowed to adhere for 4 hours at room temperature in a humidified chamber. After 4 hours, the dishes were flooded with Liebovitz medium (L-15, Gibco, 11415064) supplemented with 10% FBS, 10 mM glucose, 10 mM phosphate buffer, and 50 U/mL Pen/Strep. The cultures were then returned to the humidified chamber and left undisturbed for 24 hours.

### Electrophysiology with Mouse Nodose Cultures

After approximately 24 hours in culture, recordings were done using a Multiclamp 700B (Molecular Devices) patch clamp amplifier and PClamp 9 acquisition software (Molecular Devices, San Jose, CA) at room temperature. Sampling of recordings was carried out at 20 KHz and filtered at 3 KHz using Digidata 1550B (Molecular Devices). Pipettes were pulled from glass capillaries (Outer diameter 1.5 mm; Inner diameter 1.1 mm) (Sutter Instruments, Novato, CA, BF150-110-10) using a PC-100 puller (Narashige, Amityville, NY) and heat-polished to a resistance of 3-5 MΩ using a microforge (MF-83, Narashige). Series resistance was generally 7 MΩ and was compensated up to 60 %. The neurons included in the final analysis had a resting membrane potential (RMP) more negative than -40 mV. After the whole-cell configuration was achieved, the RMP of all the cells was observed for about 3-4 minutes before recording was started.

All recordings were carried out at the RMP of the cells when first patched. APs were elicited from the cells by injecting slow depolarizing currents from 100 to 700 pA with a (Δ) of 200 pA throughout 1 second to mimic slow depolarization. The pipette solution contained the following (in mM): 120 K-gluconate, 6 KCl, 4 Adenosine Triphosphate (ATP)-Mg, 0.3 GTP-Na, 0.1 EGTA, 10 HEPES, and 10 phosphocreatine, pH 7.3 (with N-methylglucamine), and osmolarity was ∼290 mOsm. The external solution contained the following (in mM): 135 NaCl, 2 CaCl_2_, 1 MgCl_2_, 5 KCl, 10 glucose, and 10 HEPES, pH 7.4 (adjusted with N-methylglucamine), and osmolarity was adjusted to ∼315 mOsm with sucrose. The external bath solution containing 0.8% DMSO was used as a vehicle for all the experiments. Ramp current-evoked firing of nodose cells was initially recorded following a 2-minute perfusion of the patched cells with vehicle solution, which served as the baseline firing measurement. The cells were washed out for 5 minutes with the external bath solution (not containing 0.8% DMSO). After the 5 minutes of washout, the cells were again treated for 2 minutes with the test compounds (Mtb extract, Mtb Δ*stf0* extract, Mtb Δ*stf0::stf0* extract). Ramp-evoked firing of nodose neurons was subsequently recorded. The junction potential was calculated using the clampex function and was found to be 4.9 mV. A spike was considered an action potential only if it passed the voltage of 0 mV. The action potential properties reported such as amplitude, rheobase and latency to first spike were determined for the first action potential evoked.

### Human Dorsal Root Ganglia Cultures

Human DRG (hDRGs) tissues were recovered according to procedures described previously (Shiers et. al., 2022). The Institutional Review Board at the University of Texas at Dallas approved all the procedures and protocols regarding procurement and ethical regulations through protocol No. Legacy-MR-15-237. DRGs were procured from organ donors through Southwest Transplant Alliance (STA), a Texas organ procurement organization (OPO). STA obtained the informed consent for research tissue donation through first-person consent (driver’s license or legally binding document) or from the next of kin for the donor. Ethical oversight of OPOs is maintained by federal agencies such as the Health Resources and Services Administration (HRSA), the Centers for Medicare and Medicaid Services (CMS), and the United Network for Organ Sharing (UNOS). These agencies ensure that donation practices adhere to ethical standards, emphasizing informed consent and respect for donor autonomy.

Tissues were first isolated in the operating room at the site of organ recovery, hDRGs were maintained in artificial cerebrospinal fluid (ACSF) consisting of 95 mM NMDG, 2.5 mM KCL, 1.25 mM NaH_2_PO_4_, 30 mM NaHCO_3_, 20 mM HEPES, 25 mM Glucose, 5 mM Ascorbic acid, 2 mM Thiourea, 3 mM Sodium pyruvate, 10 mM MgSO_4_, 0.5 mM CaCl_2_, 12 mM N-acetylcysteine kept on ice. After returning to the lab, the tissue was transferred to a fresh ACSF solution and kept on ice. The tissue was carefully trimmed and cut into small pieces to enable uniform digestion and then transferred to a tube containing 1 mL of 10 mg/mL Stemxyme 1, Collagenase/Neutral Protease (Dispase, Worthington Biochemical, #LS004107), and the tube was placed in a sideways-shaking water bath. The tissue was allowed to digest for 3-4 hours with regular trituration with fire-polished glass pipettes every hour. After sufficient digestion of the tissue, it was passed through a 100 µm cell strainer (Corning, 352360). The resulting suspension was then transferred carefully using a pipette to a density gradient made of 10 % BSA (Biopharm, 71-015-025) and centrifuged at 900 x g (9 acceleration, 7 deceleration) for 5 minutes. The resulting supernatant was carefully discarded, and care was taken not to disturb the pellet. The pellet was then resuspended in media containing BrainPhys media (Stemcell technologies, Vancouver, BC #05790), 1 % Pen/Strep (Thermo, #15070063), 1 % GlutaMAX (United States Biological, #235242), 2 % NeuroCult SM1 (Stemcell technologies, #05711), 1 % N-2 Supplement (Thermo, #17502048), 2 % HyClone™ FBS (Thermo, #SH3008803IR), 1:1000 FRD+U (Sigma-Aldrich, 856657), and 10 ng/mL human βNGF (R&D systems, 256-GF). Cell density was determined by plating a 5 µL aliquot of the resuspended pellet onto a dish and counting the number of cells using a hemocytometer slide. The cells were then plated on PDL-coated coverslips at a density of 150-200 cells per coverslip and incubated at 37 °C (5 % CO_2_) for 3 hours before flooding with the same media described above. The donor demographics are listed in **Table 1**.

**Table 1:** Donor demographics for the hDRG cultures used in this study.

| Donor No. | Sex | Age | DRG levels | Cultures used for |
| --- | --- | --- | --- | --- |
| 1 | F | 45 | T11* | Ca<sup>2+</sup> Imaging |
| 2 | M | 23 | L5<sup>#</sup> | Ca<sup>2+</sup> Imaging |
| 3 | M | 53 | L4<sup>#</sup> | Ca<sup>2+</sup> Imaging |
| 4 | M | 23 | T10*, T12* and L1<sup>#</sup> | Ca<sup>2+</sup> Imaging |
| 5 | F | 45 | L1<sup>#</sup> | Ca<sup>2+</sup> Imaging |
| 6 | F | 63 | L5<sup>#</sup> | Ca<sup>2+</sup> Imaging |
| 7 | F | 35 | T10* | Ca<sup>2+</sup> Imaging |

\*Thoracic; Lumbar

### hDRG Ca^2+^ Imaging

hDRG Ca^2+^ imaging was conducted on day 4 post-plating, with media changes performed every other day. Fluo-4 dye (Invitrogen, F14201) was prepared by dissolving it in 82 µL of HBSS and 9.12 µL of Pluronic F-127 (Invitrogen, F1225), and subsequently used for imaging. Cells were loaded with dye by adding 5 µL of the dye solution to 1 mL of HBSS and incubating at 37 °C (5 % CO_2_) for 1 hour. After the hour incubation, the coverslips were transferred to a perfusion slide. External bath containing 125 mM NaCl, 4.2 mM KCl, 29 mM NaHCO_3_, 20 mM Glucose, 2 mM MgSO_4_, 1 mM CaCl_2_, 2.5 mM KCl, 1.25 mM NaH_2_PO_4_, pH 7.4 with NMDG and osmolality of 300-305 mOsm was added and perfused for 1 min before the start of imaging. For experiments done without Ca^2+^, the external bath was modified to contain 125 mM NaCl, 4.2 mM KCl, 29 mM NaHCO_3_, 20 mM Glucose, 3 mM MgSO_4_, 2.5 mM KCl, 1.25 mM NaH_2_PO_4_, pH 7.4 with NMDG and osmolality of 300-305 mOsm.

Cells were subjected to treatments like those described for mouse nodose neurons, with the exception that the exposure duration for each tested compound was approximately 150 seconds. For experiments involving YM, with or without Ca^2+^ in the external bath, cells were incubated in 100 µM YM before imaging. The data were normalized by subtracting the treatment periods from the entire 50 second baseline as the measurements obtained with Flo-4 dye were not ratiometric. Data analysis was performed following the same procedures used for mouse nodose cells.

### Statistical Analysis

GraphPad Prism software was used to perform statistical analysis between groups. Statistical analysis between multiple groups was performed using two-way ANOVA and multiple comparisons with a Bonferroni post-hoc test. Differences between groups were considered significant at *p* < 0.05. All electrophysiology traces were analyzed using Clampfit 10 to determine rheobase, latency to first spike, and amplitude. No sex-specific effects were observed in response to any of the treatments, and both male and female tissue samples were used to obtain the data.

## Results

### Mtb extract increases the firing rate of Mouse nodose neurons

We previously demonstrated that Mtb extract induces Ca^2+^ signaling in mouse and human DRG neurons, and in mouse nodose neurons, was dependent on the presence of SL-1 in the Mtb extract (Ruhl et al., 2020). However, whether Ca^2+^ signaling in nodose neurons in response to Mtb extract sensitizes for action potential generation is unknown. To address this knowledge gap, we tested the impact of Mtb extract and SL-1 on modulation of evoked action potentials in cultured mouse nodose neurons. The general schematic for the treatment and recording of ramps is shown in **Figures 1A** and **B**. The groups evaluated consisted of bath (0.8% DMSO) vs. Mtb extract (0.4 mg/ml, n = 8), bath (0.8% DMSO) vs. Mtb Δ*stf0* extract (0.4mg/ml, n=8), and bath (0.8% DMSO, n = 8) vs. Mtb Δ*stf0*::*stf0* extract (0.4 mg/ml, n = 11). Cells included in the final analysis had a similar mean diameter in µm in each of the treatment groups tested: Mtb extract vs. bath (0.8% DMSO) 24.59 ± 0.86; Mtb Δ*stf0* extract vs. bath (0.8% DMSO) 23.51 ± 0.7; and Mtb Δ*stf0*::*stf0* extract vs. bath (0.8% DMSO) 24.39 ± 0.9. APs were elicited in these cells by injecting slow ramp currents (see methods). An increase in the AP firing was observed in nodose neurons in response to a 2-minute treatment with Mtb extract (**Figures 2A** and **B**). In contrast, treatment with the Mtb Δ*stf0* extract lacking SL-1 (Ruhl et al., 2020) did not elicit increased action potential firing in response to injection of ramp currents (**Figures 2C** and **D**). Finally, exposure of neurons to the Mtb Δ*stf0*::*stf0* extract containing SL-1 restored the increased action potential firing in response to injected ramp current in nodose neurons (**Figures 2E** and **F**, ramps for each protocol shown in **Figure 2G**).

**Figure 1:**
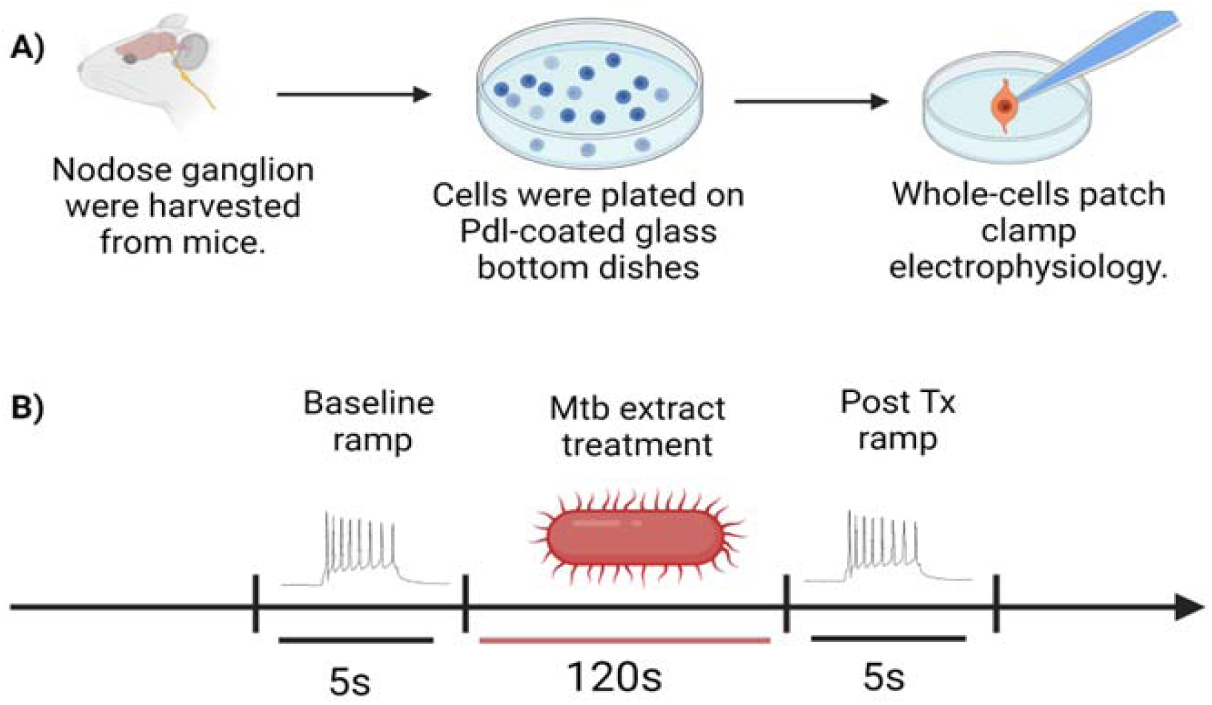
Schematics showing **A**) the culture process of nodose neurons from mice, **B)** the process for recording ramp current-evoked AP generation from mouse nodose neurons using whole-cell patch clamp electrophysiology of bath (vehicle) vs. Mycobacterium*-*derived extracts.

**Figure 2:**
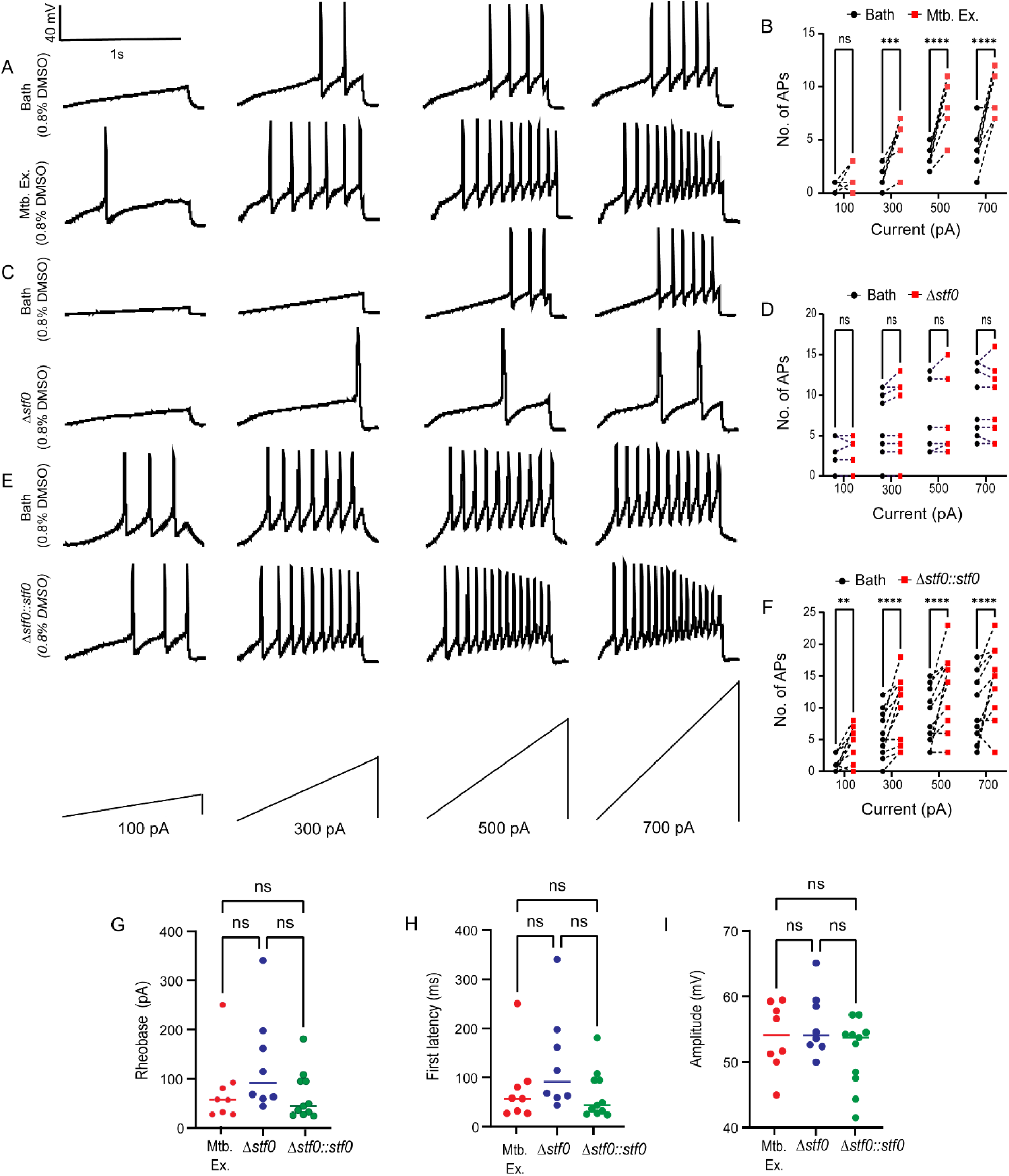
Total lipid extract from Mtb can induce increased AP generation in mouse nodose neurons when stimulated with ramp currents. (A) Representative traces of showing ramp-induced firing at 100, 300, 500, and 700 pA in mouse nodose neurons before and after treatment with Mtb extract. (B) Comparison between the number of APs fired by mouse nodose neurons when treated with bath and Mtb extract (0.8% DMSO, n = 8). (C) Representative traces showing ramp-induced firing in mouse nodose neurons before and after treatment with Mtb Δ*stf0* extract at 100, 300, 500, and 700 pA. (D) Number of APs fired by mouse nodose neurons when treated with bath and Mtb Δ*stf0* extract (0.8% DMSO, n = 8). (E) Representative traces of showing ramp-induced firing in mouse nodose neurons before and after treatment with Mtb Δ*stf0*::*stf0* extract at 100, 300, 500, and 700 pA. (F) Comparative analysis of APs fired in response to treatment with bath and Mtb Δ*stf0*::*stf0* extract (0.8% DMSO, n = 11).(G, H, and I) Graphs representing comparative analyses of the rheobase, latency to first action potential, and amplitude in response to Mtb extract, Mtb Δ*stf0*, and Mtb Δ*stf0*::*stf0* extracts. p-values were calculated using two-way ANOVA with correction for multiple comparisons by Bonferroni post hoc test.

We also assessed action potential related membrane properties such as rheobase, latency to first spike, and action potential amplitude using APs elicited by a 500-pA ramp current injection (**Figures 2H, I, J**). This current amplitude was selected because all cells across treatment groups consistently fired APs at this level. None of the action potential properties were altered across all treatment groups. These findings suggest that Mtb and Mtb Δ*stf0*::*stf0* extracts enhance the firing rate of nodose neurons in response to slow ramp current injections, without altering action potential properties such as rheobase, latency to first spike, or action potential amplitude. This modulation of neuronal excitability is dependent on SL-1, as evidenced by the unchanged firing rates observed in neurons treated with the Mtb Δ*stf0* extract devoid of SL-1.

### YM inhibits Ca^2+^ responses elicited in response to Mtb extract and SL-1 in mouse nodose neurons

Previous studies have shown that Mtb extract and purified SL-1 induce Ca^2+^ signaling in mouse nodose neurons with a Ca^2+^ spike that reaches its peak 45 seconds post-addition of both compounds (Ruhl et al., 2020). This slow Ca^2+^ response is consistent with a Gαq/11 coupled GPCR activating release of Ca^2+^ from intracellular stores through the phospholipase C enzyme (PLC)/inositol trisphosphate second messenger (IP3)/diacylglycerol second messenger (DAG) pathway (Dhyani et al., 2020; Predescu et al., 2019). Elucidating the source of Ca^2+^ mobilized by SL-1 could provide valuable insights into its potential target receptors and underlying mechanisms of action.

Ca^2+^ signaling was first tested with the presence of Ca^2+^ in the external bath solution. Cells were treated with vehicle and Mtb extract, followed by Cap to assess their response to the established TRPV1 agonist (**Figures 3A** and **B**). Mtb extract and Cap activated nodose neurons as indicated by an increase in Ca^2+^ levels in these cells. One third (29.9 %) of the cells tested responded to Mtb extract and Cap (**Figure 3C**) and all Mtb extract responsive cells were also responsive to Cap. Non-neuronal cells were not responsive to Mtb extract. To determine if the Ca^2+^ release triggered by exposure to Mtb extract was Gαq/11 mediated, we pretreated the nodose neurons with YM (100 μM) for 15 minutes before imaging (Trent et al., 2024). Pretreatment with YM effectively inhibited the Mtb extract induced Ca^2+^ signal in nodose neurons, suggesting that the response was Gαq/11 dependent (**Figures 3D** and **E**). The area under the curve analysis (AUC) showed that the Ca^2+^ spike in cells treated with Mtb extract alone was significantly higher than either vehicle treatment or in cells pretreated with YM prior to exposure to Mtb extract (**Figure 3F**).

**Figure 3:**
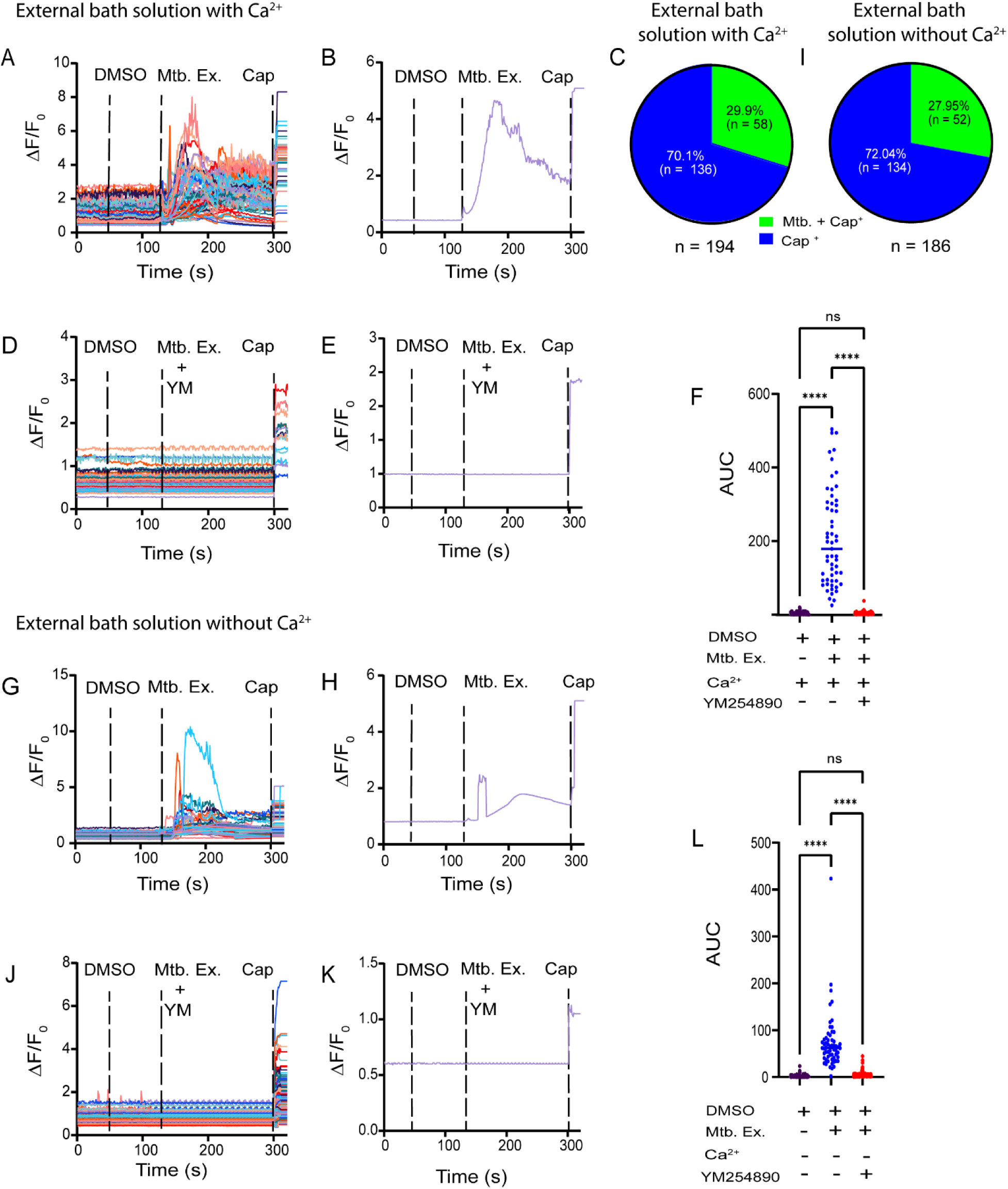
Mtb extract activates mouse nodose neurons in a YM reversible fashion that does not depend on external Ca^2+^. (A) Traces demonstrating Ca^2+^ increase from mouse nodose neurons treated with Mtb extract and Cap with Ca^2+^ in the external bath solution. (B) Representative trace for response to Mtb extract and Cap plotted against time (seconds). (D) Nodose neurons treated with YM (100 µM) can inhibit the release of Ca^2+^ when treated with Mtb extract. and Cap with Ca^2+^ in the external bath solution. (E) Single representative trace showing response to Mtb extract plotted against time (seconds) when cells are pretreated with YM. (G) Combined traces showing the release of Ca^2+^ from nodose neurons activated by Mtb extract and Cap in the absence of Ca^2+^ in the external bath solution. (H) Single representative trace showing response to Mtb extract in the absence of Ca^2+^ in the external bath solution. (J) Combined traces showing no Ca^2+^ release in nodose neurons treated with YM when activated by Mtb extract in the absence of Ca^2+^ in the external bath solution. (K) Single representative trace showing no response to Mtb extract in the absence of Ca^2+^ in the external bath solution. (C,I) Pie chart showing the percentage of nodose neurons responsive to both Mtb extract and Cap, and Cap alone in the presence or absence of Ca^2+^ in the external bath solution. (F, L) AUC analysis for cell responses to vehicle (DMSO), Mtb extract in the presence or absence of YM, and Ca^2+^ in the external bath solution using two-way ANOVA multiple comparison with a Bonferroni post-hoc test.

Having established that Mtb extract induced Ca^2+^ signaling in nodose neurons is inhibited by pretreatment with YM, we next aimed to identify the specific source of Ca^2+^. We removed Ca^2+^ from the external bath and tested for neuronal Ca^2+^ levels after exposure to Mtb extract (**Figures 3G** and **H**). Approximately 50 seconds after the addition of Mtb extract, a pronounced Ca^2+^ signal was observed in many nodose neurons, which were again all Cap sensitive (Ca^2+^ in external solution was replenished prior to Cap application). Again, about a quarter (27.95%) of the cells responded to Mtb extract and Cap, a proportion comparable to the responses observed in the presence of extracellular Ca^2+^ (**Figure 3I**). We further evaluated whether Mtb extract evoked Ca^2+^ signaling could be inhibited using YM in a Ca^2+^ free external bath solution (**Figures 3J** and **K**). As with the experiment conducted in the presence of extracellular Ca^2+^, YM treatment prevented the Mtb extract induced increase in intracellular Ca^2+^ (**Figure 3L**).

We next assessed the impact of pure SL-1 (1 µM) on mouse nodose neurons. SL-1 alone led to Ca^2+^ signaling in a substantial subset of nodose neurons that also responded to Cap (**Figures 4A** and **B**). This experiment was carried out in the presence of Ca^2+^ in the external bath. Of the cells imaged, 31.6 % responded to both SL-1 and Cap (**Figure 4C**) and no cells responded to SL-1 and not Cap. To further demonstrate that the Ca^2+^ signal elicited in the neuronal cells in response to SL-1 treatment was Gαq/11 mediated, we pretreated cultures with YM as before. In cells pretreated with YM, a complete inhibition of Ca^2+^ signaling was observed (**Figures 4D** and **E**). AUC analysis showed that Ca^2+^ increases in cells treated with SL-1 was significantly higher than both vehicle only and cells exposed to YM and SL-1 (**Figure 4F**).

**Figure 4:**
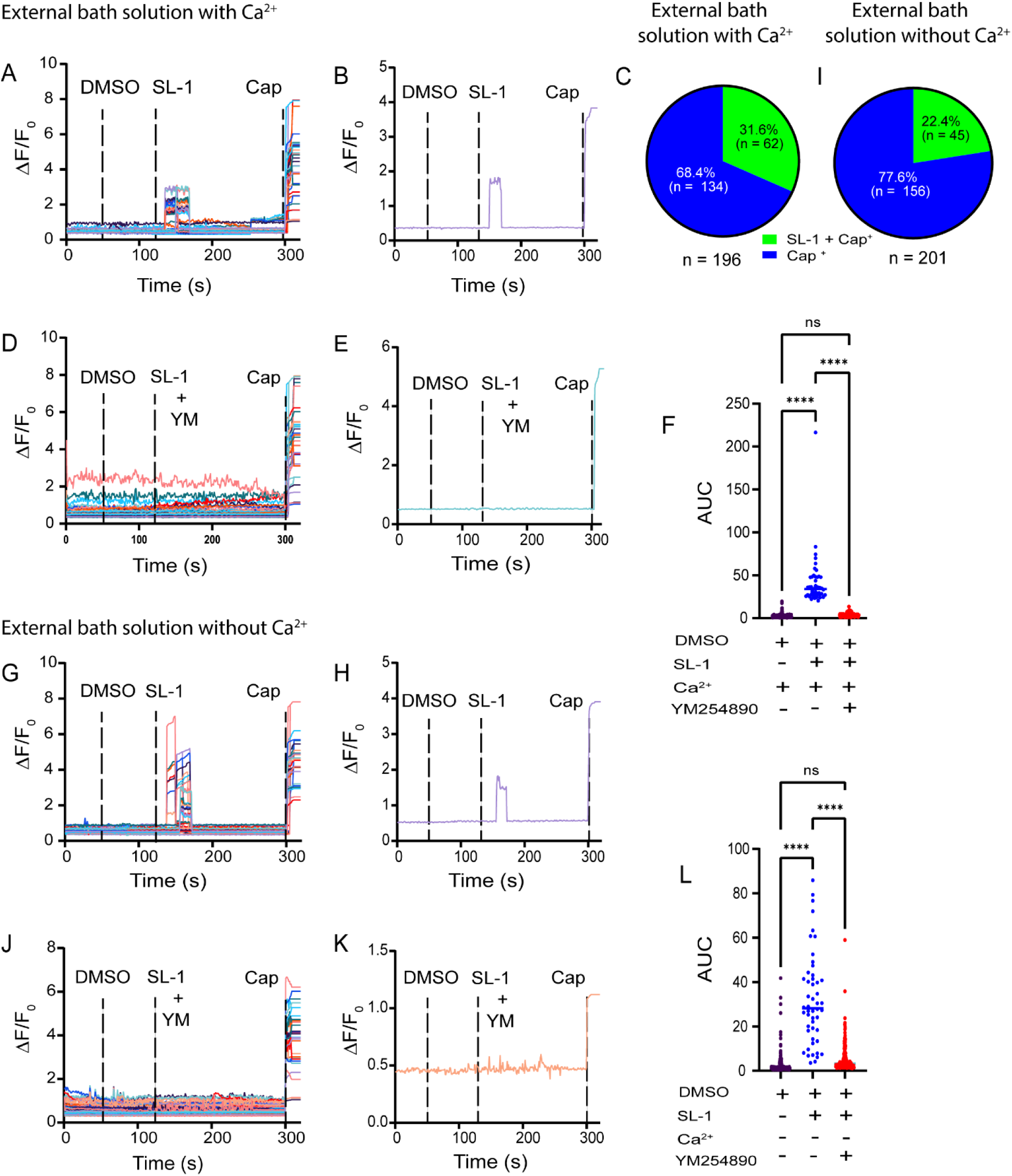
SL-1 induced Ca^2+^ signaling in mouse nodose neurons in a YM reversible fashion that does not depend on external Ca^2+^. (A) Traces demonstrating the Ca^2+^ release from nodose neurons treated with SL-1 and Cap with Ca^2+^ in the external bath solution. (B) Representative trace for response to SL-1 and Cap plotted against time (seconds). (D) Nodose neurons treated with 100 µM of YM can inhibit the release of Ca^2+^ when treated with SL-1 and Cap with Ca^2+^ in the external bath solution. (E) A single representative trace showing a response to SL-1, plotted against time (seconds). (G) Combined traces showing the release of Ca^2+^ from nodose neurons activated by SL-1 and Cap in the absence of Ca^2+^ in the external bath solution. (H) A representative trace showing a response to SL-1 in the absence of Ca^2+^ in the external bath solution, plotted against time (seconds). (J) Combined traces showing no Ca^2+^ release in nodose neurons treated with YM and SL-1 in the absence of Ca^2+^ in the external bath solution. (K) Single representative trace showing no response to SL-1 in the absence of Ca^2+^ in the external bath solution, plotted against time (seconds). (C,I) Pie charts represent the percentage of nodose neurons responsive to both SL-1 and Cap, and Cap alone in the presence or absence of Ca^2+^ in the external bath solution. (F,L) Area under the curve analyses cell responses to DMSO, SL-1 in the presence or absence of YM, and Ca^2+^ in the external bath solution using two-way ANOVA multiple comparison with a Bonferroni post-hoc test.

We then tested neuronal activation with SL-1 in the absence of Ca^2+^ in the external bath (**Figures 4G** and **H**). We observed that SL-1 triggered a Ca^2+^ signal in these cells approximately 45 seconds post addition of the compound in the absence of external Ca^2+^. Of the cells tested, 22.4 % of the cells responded to both SL-1 and Cap, a proportion slightly lower when compared to the responses observed in the presence of external Ca^2+^ (**Figure 4I**). We then assessed if SL-1-mediated responses in mouse nodose neurons can be inhibited by YM in a Ca^2+^ free external bath solution (**Figures 4J** and **K**). SL-1 failed to elicit a Ca^2+^ release under these conditions, similar to observations with Mtb extract (**Figures 3L** and **4L**). Taken together, we conclude that Ca^2+^ mobilized by SL-1 originates from intracellular stores in a mechanism dependent on Gαq/11 signaling.

In a set of control experiments, we evaluated the Mtb Δ*stf0* extract, which lacks SL-1, under conditions both with and without extracellular Ca^2+^ and in the presence or absence of the Gαq/11 inhibitor YM (**Figures 5A-H**). Mtb Δ*stf0* extract failed to elicit Ca^2+^ release under any of the conditions. These results further indicate that SL-1 is the component in Mtb extract that drives Gαq/11 dependent signaling leading to release of intracellular Ca^2+^ stores in mouse nodose neurons.

**Figure 5:**
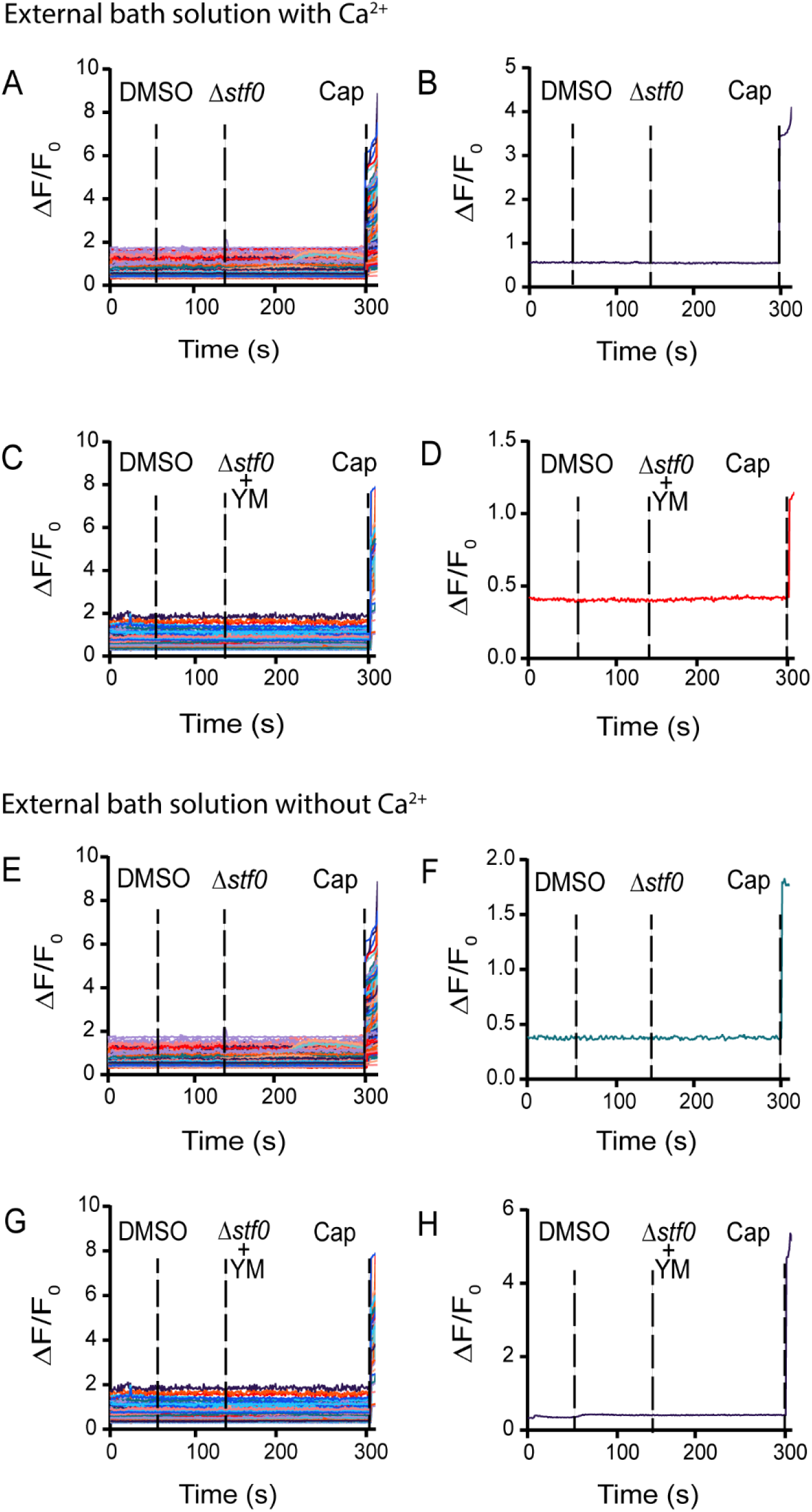
Extracts derived from Δ*stf0* do not trigger the release of Ca^2+^ in mouse nodose neurons in the presence or absence of extracellular Ca^2+^. (A, C) Combined traces from mouse nodose neurons causing no Ca^2+^ release when treated with Δ*stf0* and YM in the presence of extracellular Ca^2+^. (B, D) Single traces from mouse nodose neurons in response to Δ*stf0* and YM treatment in the presence of extracellular Ca^2+^, plotted against time (seconds). (E, G) Combined traces from nodose neurons showing a lack of Ca^2+^ release when treated with Δ*stf0* and YM in the absence of extracellular Ca^2+^. (F, H) Single traces from mouse nodose neurons in response to Δ*stf0* and YM treatment in the absence of extracellular Ca^2+^, plotted against time (seconds).

### YM inhibits Ca^2+^ responses elicited in response to Mtb extract and SL-1 in hDRG neurons

Previous studies have shown that Mtb extract and SL-1 lead to Ca^2+^ signaling in hDRG neurons, but these experiments were done using a limited number of neurons, and the underlying mechanisms driving this signaling were not determined (Ruhl et al., 2020). Since Mtb primarily infects humans, we wanted to validate the mouse nodose Ca^2+^ imaging data using a human neuronal system. We first tested hDRGs for activation of Ca^2+^ signaling with Mtb extract and SL-1 in the presence of Ca^2+^ in the external bath solution (**Figures 6A-B** and **C-D**). We observe that Mtb extract produced a Ca^2+^ signal within approximately 30 seconds of cellular exposure to the extract. When hDRG neurons were tested with SL-1, they showed a Ca^2+^ signal approximately 50 seconds post addition. A majority of cells (59 %) responded to both SL-1 and Cap and 49 % responded to both Mtb extract and Cap. Like mouse neurons, all hDRG neurons that responded to SL-1 or Mtb extract also responded to Cap, highlighting their nociceptive nature.

**Figure 6:**
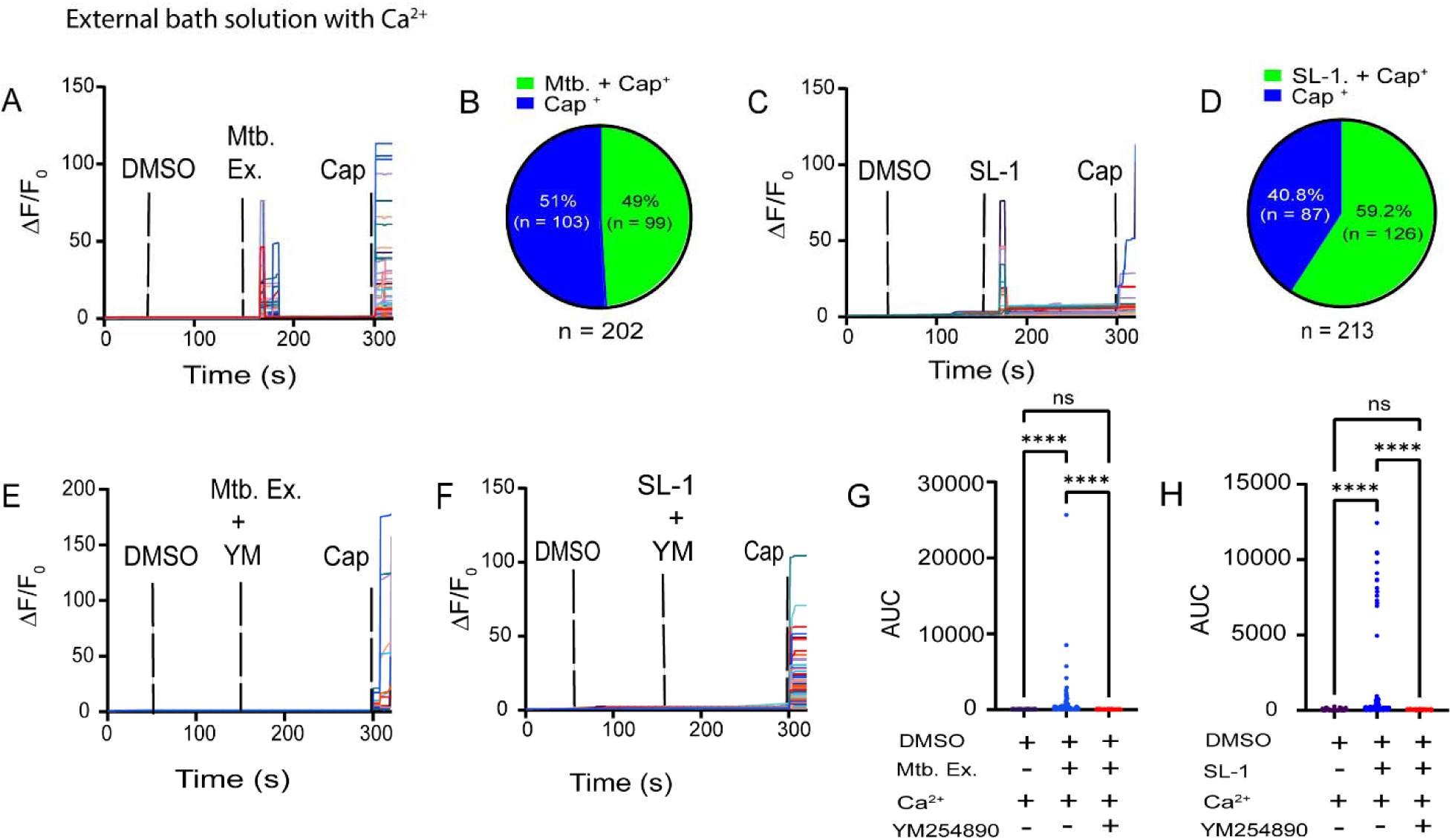
SL-1 and Mtb extract activate Ca^2+^ signaling hDRG neurons in a YM reversible fashion. (A) Combined traces demonstrating the Ca^2+^ release from hDRG neurons treated with Mtb extract and Cap with Ca^2+^ in the external bath solution, plotted against time (seconds). (B) Pie chart showing the number of cells responsive to Mtb extract and Cap, and Cap alone, in the presence of external Ca^2+^ in the external bath solution. (C) Traces demonstrating the release of Ca^2+^ from hDRG neurons treated with SL-1 and Cap, with Ca^2+^ in the external bath solution, plotted against time (seconds). (D) Pie chart showing percentage of cells responsive to SL-1 and Cap, and Cap alone in the presence of external Ca^2+^ in the bath solution. (E) Combined traces showing hDRG treated with 100 µM of YM showed inhibition of Ca^2+^ release when treated with Mtb extract and Cap, with Ca^2+^ in the external bath. (F) Combined traces showing no Ca^2+^ release in hDRG neurons treated with YM and SL-1, in the presence of external Ca^2+^ in the external bath solution. (G) AUC representing cell responses to DMSO, Mtb extract, in the presence and absence of YM, with Ca^2+^ present in the external bath solution. (H) AUC analysis for cell responses to DMSO, SL-1 in the presence and absence of YM, and presence of Ca^2+^ in the external bath solution.

Next, we evaluated whether the Ca^2+^ signaling elicited by both Mtb extract and SL-1 could be inhibited by YM (**Figures 6E** and **F**). No Ca^2+^ signal was elicited by either Mtb extract or SL-1 when cells were pretreated with YM. This is further highlighted in both cases the significant difference in AUC between the treatment groups (**Figures 6G** and **H**).

We next evaluated the source of the Ca^2+^ observed in response to both Mtb extract and SL-1. To do this, we evaluated the impact of Mtb extract and SL-1 on hDRGs in a Ca^2+^ free external bath solution (**Figures 7A-B** and **7C-D**). We observed that Mtb extract produced a Ca^2+^ signal in hDRG cells soon after its addition, while in the case of SL-1, there was a latency of approximately 50 seconds. 32 % of hDRG neurons responded to both Mtb extract and Cap, while 25 % of the cells responded to both SL-1 and Cap. To further investigate whether the Ca^2+^ release induced by Mtb extract and SL-1 in hDRG neurons was mediated by Gαq/11 signaling, we pretreated neurons with YM in a Ca^2+^ free external bath solution (**Figures 7E** and **F**). No Ca^2+^ signaling was observed in response to either Mtb extract or SL-1 in hDRG neurons. AUC analysis showed a significant difference in Ca^2+^ signaling between all treatment groups described above (**Figures 7G** and **H**).

**Figure 7:**
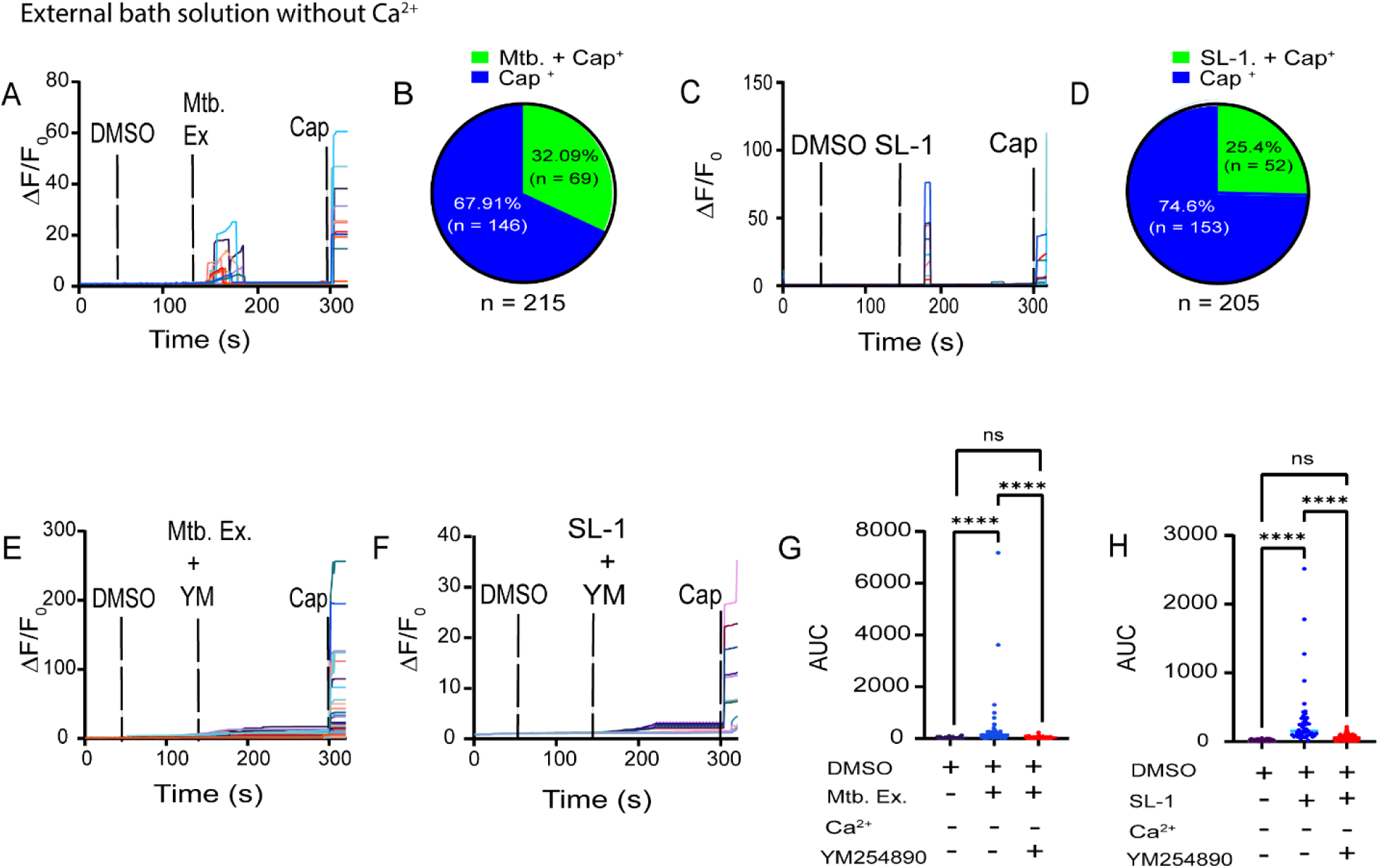
Mtb extract and SL-1 induce Ca^2+^ signaling in hDRG neurons in a YM reversible fashion in the absence of extracellular Ca^2+^. (A) Combined traces showing release of Ca^2+^ from hDRG neurons as activated by Mtb extract and Cap, in the absence of Ca^2+^ in the external bath solution, plotted against time (seconds). (B) Pie chart represents the number of cells responsive to Mtb extract and Cap, and Cap alone in the absence of external Ca^2+^ in the bath. (C) Combined traces showing release of Ca^2+^ in hDRG neurons, when activated by SL-1 in the absence of external Ca^2+^, in the external bath solution, plotted against time(seconds). (D) The pie chart represents the number of cells responsive to SL-1 and Cap, and Cap alone, in the absence of external Ca^2+^ in the bath solution. (E) Combined traces showing release of Ca^2+^ from hDRG neurons treated with YM and activated by Mtb extract and Cap, in the absence of Ca^2+^, in the external bath solution, plotted against time (seconds). (F) Combined traces showing no Ca^2+^ release in hDRG neurons treated with YM and SL-1 in the absence of external Ca^2+^, in the external bath solution, plotted against time (seconds). (G) AUC of cell responses to DMSO, Mtb extract in the presence and absence of YM, and absence of Ca^2+^ in the external bath solution. Data analyzed using two-way ANOVA, multiple comparisons with Bonferroni post-hoc test. (H) AUC analysis for the cell responses to DMSO, SL-1 in the presence and absence of YM, and absence of Ca^2+^ in the external bath solution, using two-way ANOVA multiple comparisons with a Bonferroni post-hoc test.

Finally, as an additional control, we evaluated the impact of the Mtb Δ*stf0* extract on hDRG neurons under conditions both with and without extracellular Ca^2+^ and in the presence or absence of YM (**Figures 8 A-E**). In all conditions the Mtb Δ*stf0* extract failed to elicit a Ca^2+^ response in hDRG neurons.

**Figure 8:**
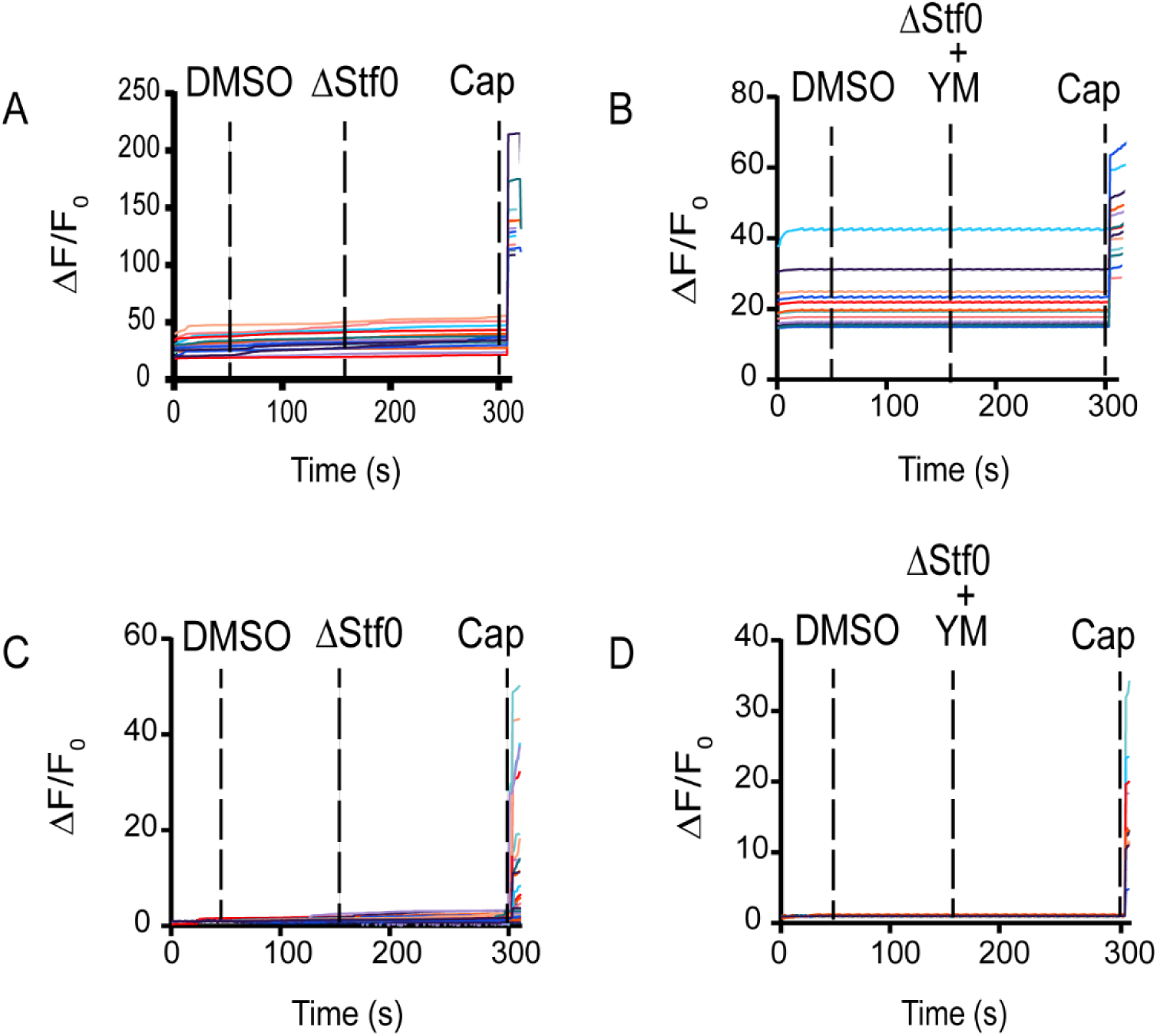
Mtb Δ*stf0* extract does not trigger intracellular Ca^2+^ in hDRG in the presence or absence of extracellular Ca^2+^. (A) Combined traces from hDRG neurons in response to treatment with Mtb Δ*stf0* extract in the presence of extracellular Ca^2+^. (B) Combined traces of hDRG neurons treated with 100 µM YM and Mtb Δ*stf0* extract in the presence of extracellular Ca^2+^. (C) Combined traces of hDRG neurons treated with Mtb Δ*stf0* extract in the absence of extracellular Ca^2+^. (D) Combined traces of hDRG neurons treated with 100 µM YM and Mtb Δ*stf0* extract in the absence of extracellular Ca^2+^.

## Discussion

In this study, we expanded on our previous findings that both Mtb extract and SL-1, a glycolipid produced by Mtb, elicit Ca^2+^ responses in mouse nodose and hDRG neurons, with SL-1 identified as the component mediating the effect (Ruhl et al., 2020). Key findings emerging from the work described here are that Mtb extract and SL-1 activated mouse nodose and human DRG neurons through a mechanism that requires Gαq/11 signaling leading to the release of Ca^2+^ from intracellular stores. In the case of mouse nodose neurons, Mtb extracts containing SL-1 increased AP generation in response to ramp current stimulation. This increased excitability demonstrates that Mtb extract and SL-1 sensitize TRPV1^+^ nociceptors in mouse and humans, a finding that also aligns with prior research showing that bacterial metabolites and toxins can interact directly with the nervous system to modulate nervous system function (Chiu et al., 2013; Erdogan et al., 2025; Yang and Chiu, 2017).

To understand the impact of SL-1 on neuronal excitability, we employed patch clamp electrophysiology on mouse nodose neurons using a ramp protocol. We observed that Mtb extract enhanced membrane excitability to increase the firing of nodose neurons without altering action potential properties in a process dependent on the Mtb sulfotransferase stf0. Future experiments should address whether SL-1 leads to subthreshold changes in membrane potential, or generator potentials, that could increase the probability of firing an AP in the presence of other mediators (McGovern et al., 2018). Such an effect could sensitize nociceptors that initiate the cough reflex.

Release of intracellular Ca^2+^ is a hallmark of GPCR, specifically Gαq/11, signaling in cells (Bootman and Roderick, 2011; Dhyani et al., 2020; Predescu et al., 2019; Xu and Xie, 2009). Our previous studies demonstrated a latency of approximately 40 seconds before a Ca^2+^ response was elicited in response to both Mtb extract and SL-1 in mouse nodose and hDRG neurons (Ruhl et al., 2020). This finding led us to hypothesize that SL-1 acts via a metabotropic receptor signaling mechanism in mouse and human nociceptors. To formally test this hypothesis, we evaluated the effects of Mtb extract and SL-1 on Ca^2+^ release in both mouse nodose and hDRG neurons under conditions with and without external Ca^2+^. Our results indicate that both Mtb extract, and SL-1 produce a robust Ca^2+^ signal, even in a Ca^2+^ free bath. These results support the conclusion that the observed early Ca^2+^ signal is attributed to the preferential release of Ca^2+^ from internal cellular stores rather than translocation via membrane channels from the extracellular environment. Treatment with the Gαq/11 inhibitor YM likewise attenuated the Ca^2+^ responses induced by both Mtb extract and SL-1, regardless of whether Ca^2+^ was present in the extracellular bath. We noted that the response to both Mtb extract and SL-1 was blunted in the absence of extracellular Ca^2+^ suggesting that ion channel mediated mechanisms may be at play following the release of Ca^2+^ from intracellular stores, but since there was no response with YM treatment, this mechanism must require initiation of signaling that is attenuated entirely by YM pretreatment. Future investigation will assess how ion channel conductance may be modulated by SL-1 in human and mouse nociceptors. Importantly, we only observed Mtb extract- and SL-1-induced signaling in TRVP1^+^ neurons demonstrating that this action is selective for mouse and human nociceptors.

Previous studies have suggested that Gαq/11 mediated release of intracellular Ca^2+^ can influence the activity of ATP-sensitive potassium channels (Nakano et al., 2002; Oh, 2023) through the activation of the PLC-β /IP3 pathway, which leads to Ca^2+^ release from the endoplasmic reticulum. This subsequently activates phosphoinositide 3-kinase (PI3K) and results in phosphorylation of Protein kinase B (Akt) (Melchior and Frangos, 2012; Wickman and Clapham, 1995). Akt then phosphorylates and modulates ATP-sensitive potassium (K_ATP_) channels, which can increase neuronal firing without changing basic membrane properties (Bonvini and Belvisi, 2017; Nguyen et al., 2024; Sun et al., 2005; Zhuang et al., 2004). Our electrophysiology data in mouse nodose suggests that this pathway may be involved in the increased firing observed in response to SL-1 without modulation of rheobase. Furthermore, our Ca^2+^ imaging data in both mouse and human neurons strongly suggest that Gαq/11 mediated GPCR signaling drives these excitability changes via intracellular Ca^2+^ release.

A limitation of our work is that we have not identified the receptor-mediated mechanism that causes the signaling effects we have observed here. This will be a major goal of our ongoing work in understanding how TB drives interactions with nodose and DRG nociceptors. The discovery that this signaling engages a GPCR-like mechanism, is important for targeting screening efforts to identify this receptor(s). Furthermore, narrowing the mechanism to TRPV1^+^ nociceptors in both the mouse and human limits the pool of potential targets as transcriptomes of these neuronal populations are well described in both species (Bhuiyan et al., 2024; Kupari et al., 2019; Tavares-Ferreira et al., 2022; Usoskin et al., 2015; Yu et al., 2024).

Our findings demonstrate that SL-1, a component of the Mtb extract, activates Gαq/11 mediated GPCR signaling pathways, leading to intracellular Ca^2+^ release in both mouse nodose and hDRG neurons. Furthermore, electrophysiological findings in mouse nodose neurons show enhanced neuronal excitability, which may be driven downstream by K_ATP_ channels. Further work in this area should be focused on identifying specific host receptors for SL-1, which may potentially lead to the development of targeted therapies aimed at treating Mtb induced cough. These interventions could possibly assist in reducing disease transmission, particularly in regions where it remains endemic with limited access to current therapies.

## ACKNOWLEDGMENTS

The authors would like to thank all members of the Price and Dussor lab for their support and constructive feedback on the manuscript. We thank the Southwest Transplant Alliance for the recovery of tissues from organ donors, and we thank the organ donors and their families for their gift. This work was supported by NIH R01 AI158688 (M.U.S), P01 AI159402 (M.U.S) and U19NS130608 (T.J.P). Michael Shiloh would also like to acknowledge support from the Disease Oriented Clinical Scholar program at UT Southwestern.

## AUTHOR CONTRIBUTIONS

Conceptualization, D.K.N., M.U.S., and T.J.P.; Formal Analysis, D.K.N. and T.J.P.; Funding Acquisition, T.J.P and M.U.S.; Investigation, D.K.N, F.dJ.E-B, I.M.O, K.F.N, G.N. and C.R.R.; Project Administration, G.D., T.J.P. and M.U.S.; Resources, T.J.P.; Supervision, G.D., T.J.P. and M.U.S.; Writing-original draft, D.K.N. and T.J.P. Writing-Review and Editing, all authors.

